# The effector of Hippo signaling, Taz, is required for formation of the micropyle and fertilization in zebrafish

**DOI:** 10.1101/319475

**Authors:** Xiaogui Yi, Jia Yu, Chao Ma, Guoping Dong, Wenpeng Shi, Li Li, Lingfei Luo, Karuna Sampath, Hua Ruan, Honghui Huang

## Abstract

The mechanisms that ensure fertilization of eggs by a single sperm are not fully understood. In all teleosts, a channel called the ‘micropyle’ is the only route of entry for sperm to enter and fertilize the egg. The micropyle forms by penetration of the developing vitelline envelope by a single specialized follicle cell, the micropylar cell, which subsequently degenerates. The mechanisms underlying micropylar cell specification and micropyle formation are poorly understood. Here, we show that an effector of the Hippo signaling pathway, the Transcriptional co-activator with a PDZ-binding domain (Taz), plays crucial roles in micropyle formation and fertilization in zebrafish. Genome editing mutants affecting *taz* can grow to adults, however, eggs from homozygous *taz* females are not fertilized even though oocytes in mutant females are histologically normal with intact animal-vegetal polarity, complete meiosis and proper ovulation. However, *taz* mutant eggs have no micropyle. We show that Taz protein is specifically enriched from mid-oogenesis onwards in two follicle cells located at the animal pole of the oocyte, and co-localizes with the actin and tubulin cytoskeleton. Taz protein and micropylar cell are not detected in *taz* mutant ovaries. Our work identifies a novel role for the Hippo/Taz pathway in micropylar cell specification in zebrafish, and uncovers the molecular basis of micropyle formation in teleosts.

## Author summary

In many fish, sperm enters eggs through a specialized channel called the “micropyle”. The micropyle is formed by a special follicle cell, the “micropylar cell”, which sits on top of the developing egg during oogenesis, and forms the sperm entry canal. The underlying mechanisms of this process are unknown. We find that Taz, an effector of an important signaling pathway, the Hippo pathway, is specifically enriched in micropylar cells in zebrafish, and regulates formation of the micropyle. Loss of Taz function in females results in no micropylar cells, failure to form a micropyle on eggs, which are consequently, not fertilized. Our study identifies a new role for the Hippo/Taz pathway in cell fate specification in the ovary, and reveals a potential mechanism for forming the sperm entry port. Similar mechanisms might operate in other fish as well.

## Introduction

In vertebrates, fertilization occurs by two major strategies. Amniotes such as reptiles, birds and mammals, undergo copulation and internal insemination to ensure gamete fusion, and the acrosome reaction is necessary for sperm to penetrate the zona pellucida, a protective egg envelope, and fertilization occurs at any position in the egg surface [1–3]. By contrast, most teleosts undergo external fertilization. Without a recognizable acrosome reaction, teleost sperm entry relies entirely upon a specialized funnel-like structure, the micropyle, in the chorion, an acellular coat of the egg [4–6]. Morphological and physiological studies of the micropyle in a variety of different teleost species suggest that channel formation results from transformation of a special micropylar cell from mid oogenesis [7–11]. Based on the morphological criteria, the micropylar cell is considered distinct from other follicle cells (granulosa cells) over the oocyte animal pole [12, 13]. The micropylar cell gradually expands and extends constantly through the developing vitelline envelope, till the extension tip contacts with the oocyte membrane [14]. Finally, the micropylar cell degenerates, leaving a narrow canal called the ‘micropyle’ [12, 13]. Previous studies in other teleosts revealed potential drilling forces of the micropylar cell, with the aggregation and elongation of microtubes and tonofilaments in the cytoplasm bugle of micropylar cell thought to provide internal forces [14]. Two opposing rotations between the oocyte and covering follicle cell layer are thought to supply the external force for the micropylar cell [15, 16]. Studies on a zebrafish maternal-effect mutant *bucky ball* (*buc*) revealed multiple micropyles in each egg, arising from that expanded animal identity in *buc* mutant oocytes [17, 18]. Although these studies described the morphological process of micropyle formation, little is known about the molecular mechanisms underlying formation of this essential structure.

Hippo signaling plays a variety of roles in development, regeneration, tissue homeostasis, and stress response [19, 20]. The WW domain-containing transcription regulator protein 1 (*Wwtr1*) is a transcriptional co-activator with a PDZ-binding domain (Taz). Taz, together with Yes-associated protein (Yap), are downstream effectors of Hippo signaling. As a transcriptional co-activator, Taz usually binds to transcription factors, such as Teads and Smad2/3, to regulate downstream genes transcriptions to exert its functions [21]. As an oncogene, *TAZ* has been found up-regulated in many kinds of human cancers. TAZ also promotes epithelial-mesenchymal transition (EMT), migration and invasion of cancer cells, where cell morphology is altered [22]. Previous work in zebrafish (*Danio rerio*), revealed that TAZ cooperates with Yap1 to regulate the development of retinal pigment epithelium and the establishment of the posterior body shape [23, 24].

Tumors affecting two female organs, breast and ovaries, have been used extensively to study TAZ function [25, 26]. However, to date, the role of TAZ in normal oogenesis and ovary differentiation has not been investigated. In a study of *taz* function in zabrafish, we have unexpectedly found that Taz is required for the formation of micropyle during oogenesis. We show that *taz* transcripts and encoding protein are expressed maternally. When *taz* is knocked out, some homozygous *taz* mutants can survive to adulthood, and the females display a maternal-effect phenotype, producing eggs with no micropyles. Our results suggest that Taz might regulate micropylar cell specification and morphogenesis during zebrafish oogenesis.

## Results

### 1. Taz protein and *taz* transcripts are maternally expressed

To study *taz*, we first surveyed the spatiotemporal expression pattern through zebrafish development. Whole mount *in situ* hybridization (WISH) showed that *taz* transcripts are quite abundant in two-cell stage embryos, suggesting that *taz* is a maternal expression gene (Fig. 1A). This is consistent with transcriptomic datasets [27] (http://www.ensembl.org/Danio_rerio/Location/View?r=23%3A4915595–4925724). To further verify this, sectioned ovaries were examined, and we found that *taz* mRNA and Taz protein were expressed in the cortex of oocytes and the surrounding follicle cells (Fig. 1B-D). These data revealed that *taz* is a maternally expressed gene.

**Fig. 1.**
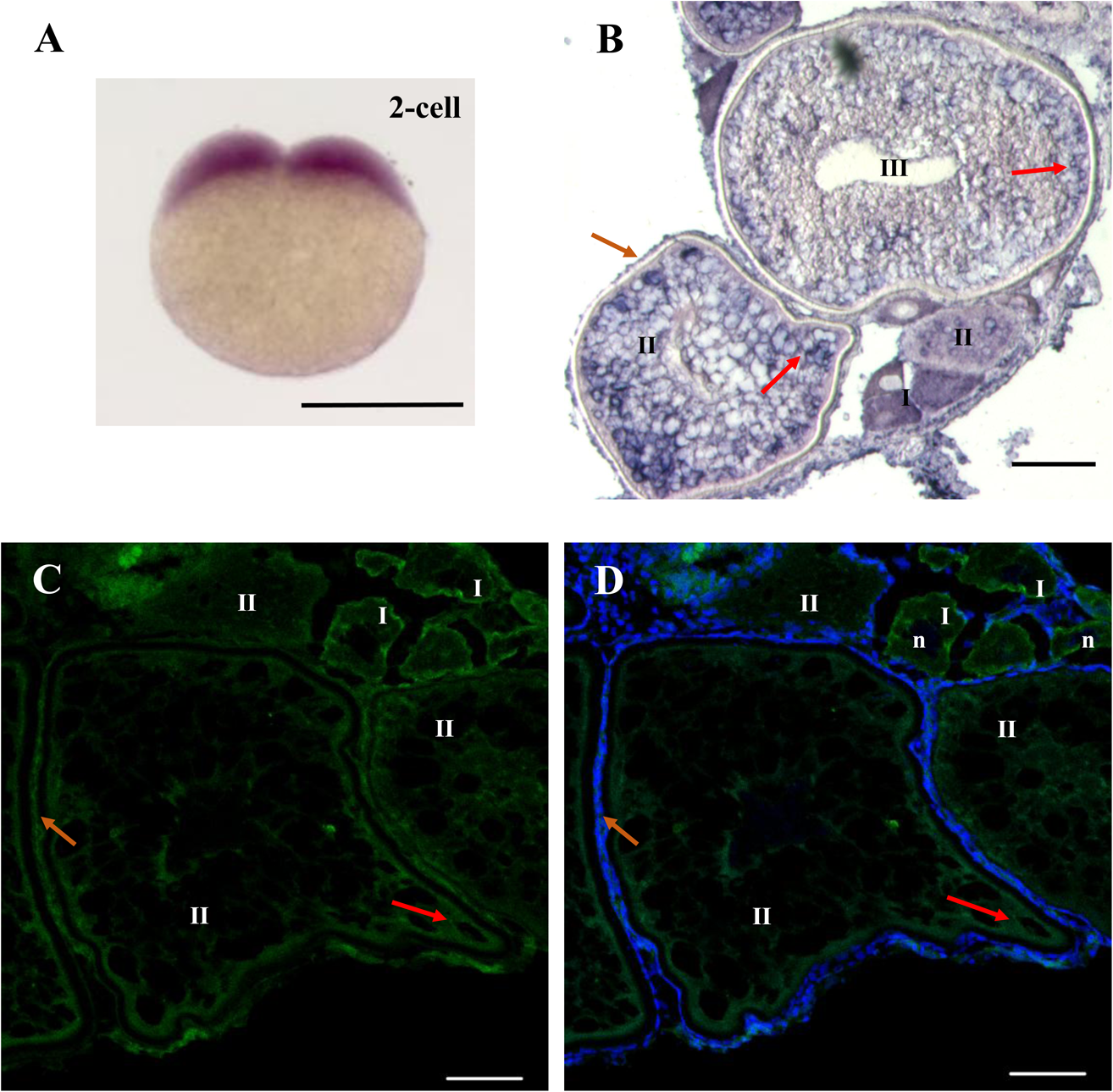
*taz* transcripts and Taz protein are expressed in oocytes and follicles cells in female zebrafish. (**A-B**) *In situ* hybridization to detect *taz* mRNA shows uniform expression in wild type embryos at the 2-cell stage (A) and in ovary sections *taz* transcripts are found in oocytes and follicle cells (B). (**C-D**) Immunofluorescence to detect Taz protein in sectioned wild type ovary. Taz protein is expressed in oocytes and follicle cells. Samples were counterstained with DAPI to detect nuclei. Oocytes at stages I, II, III are shown; n, nucleus; red arrow, the oocyte cortex; brown arrow, follicle cells. Scale bar, 500μm in A, 50μm in B, and 20μm in C-D.

### 2. Maternal Taz function is required for fertilization

To knock out *taz*, we targeted the first exon by CRISPR/Cas9 genome editing and recovered two mutant alleles, *taz^Δ10^* and *taz^Δ1^*, both of which produce mutant transcripts that encode truncated proteins with 148 and 145 amino acids, respectively (Fig. 2A-C). Importantly, no Taz protein was detected in *taz^Δ10//Δ10^* mutants (Fig. 1D). While Taz protein was found ubiquitously distributed in both oocytes and follicle cells in wild type ovaries (Fig. 2E-H), expression was totally lacking in *taz^Δ10//Δ10^* mutants (Fig. 2I-L). Taken together, these results show that the lesion in *taz^Δ10/Δ10^* results in a null mutant.

**Fig. 2.**
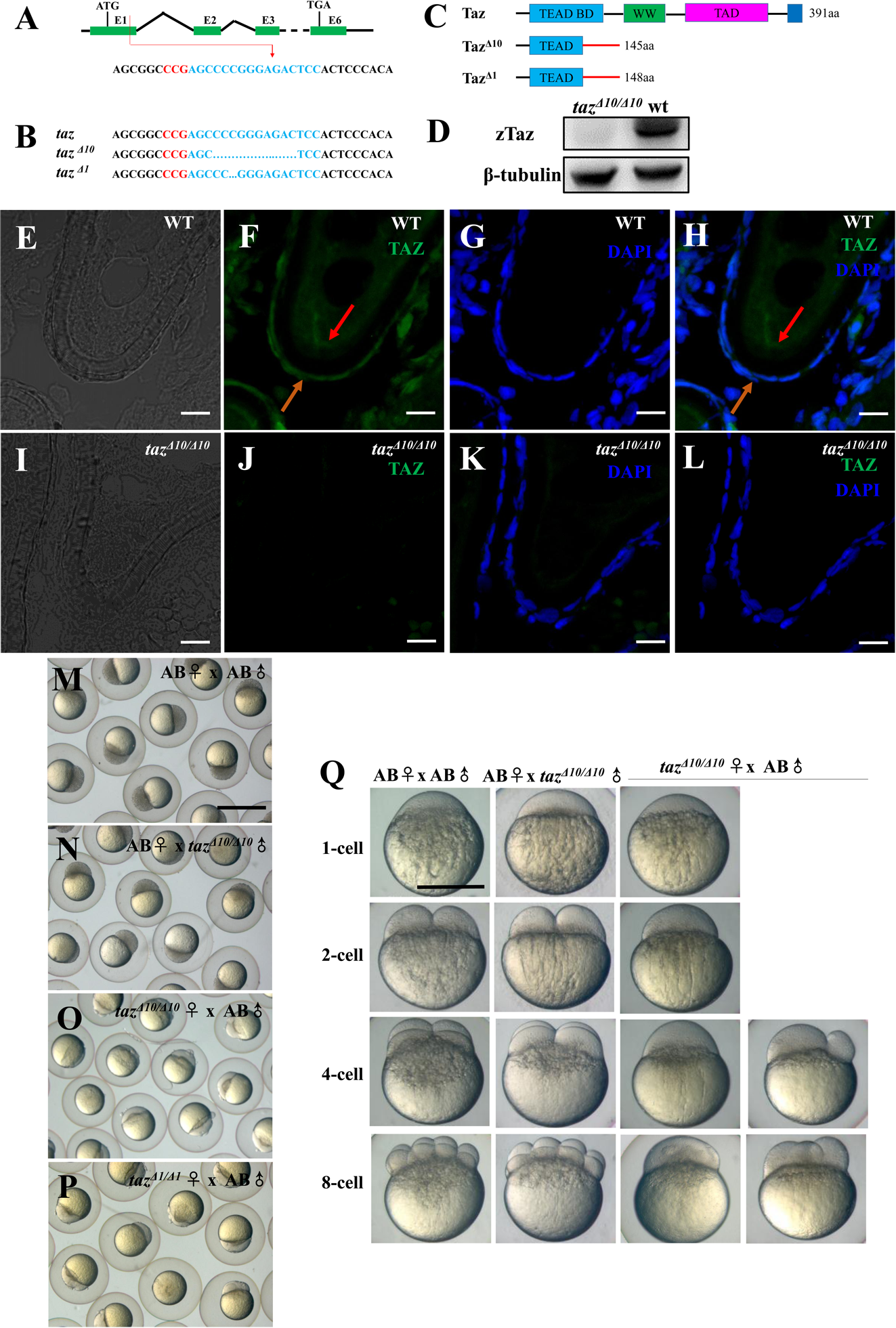
Loss of maternal Taz results in failure of eggs to develop. (**A-C**) Schematic of CRISPR/Cas9 knock-out of *taz*. The target site is located in the exon 1, the blue sequence shows the *taz* target site, whilst the red sequence is located in exon 1, and labels the PAM region (A). *taz ^Δ10^* and *taz ^Δ1^*, two mutant alleles with small deletions were obtained (B), which result in frame shifts of the *taz* open reading frame and truncated proteins (C). **(D)** Western blot analysis shows Taz protein in wild type but not in *taz^Δ10/Δ10^* mutant embryos. β-Tubulin was used as a loading control. (**E-L**) Immunofluorescence to detect Taz in sections of wild type and *taz^Δw/Δ10^* mutant ovaries. Taz protein is expressed in the cortex of the stage III oocyte and follicle cells in wild type (E-H), but is not detected in ovary sections from *taz^Δ10/Δ10^* mutants (I-L). Nuclei were counterstained with DAPI. (**M-P**) Development of embryos from crosses of wild type female with male (M), wild type female with *taz^Δ10/Δ10^* male (N), *taz^Δ10/Δ10^* female with wild type male (O) and *taz^Δ1/Δ1^* female with wild type male at 3 hpf. All embryos produced by *taz^Δ10/Δ10^* or *taz^Δ1/Δ1^* females are arrested at 1-cell stage. (**Q**) Development of embryos generated by wild type female with male, wild type female with *taz* mutant male, and *taz* mutant female with wild type male develop normally as seen at 1-cell, 2-cell, 4-cell and 8-cell stages. Most embryos produced by *taz* mutant females remain at the 1-cell stage, and a few initiate irregular cleavage planes and multiple cells at 4-cell and 8-cell stage. aa, amino acid; red arrow, the oocyte cortex; brown arrow, follicle cells. Scale bar, 10μm in E-L, 1mm in M-P, and 0.5mm in Q.

Consistent with a previous report [28], *taz^Δ10/Δ10^* mutant embryos displayed relatively normal morphology with exception of a smaller swim bladder than wild type and weak pericardial edema at 4.5 day post fertilization (dpf) (Fig. S1). Some *taz^Δ10/Δ10^* mutants obtained by intercross of heterozygotes could grow into adulthood, and the survival ratio (2.73% in 183) is much lower than the expected 25% in accordance with Mendelian segregation. Interestingly, embryos produced from mating of *taz^Δ10/Δ10^* adult females with any male arrested development at the one-cell stage, and did not survive beyond 10 hpf (Fig. 2O, 2Q), while *taz^Δ10/Δ10^* adult males were fertile (Fig. 2N, 2Q), indicating that *taz^Δ10/Δ10^* is a maternal effect mutant. The same phenomena were found in *taz^Δ1/Δ1^* allele (Fig 2P), and all subsequent studies reported in this work were done using *taz^Δ10/Δ10^* mutants.

### 3. Taz function is not required for oogenesis or oocyte polarity

To determine the basis of the failure of *taz^Δ10/Δ10^* eggs to progress beyond the one-cell stage, we examined the ovaries and oogenesis in mutant females. Compared with the same stage ovary in wild type female, the ovary of 8-month old *taz* mutant female was grossly normal in the size, tissue composition and intraperitoneal position, and there were no apparent morphological defects in the color, size and shape of oocytes (Fig. 3A, B). Histological analysis showed that all stages of oocytes (stage I to IV) were present in *taz* mutant ovaries, and had no obviously difference from that in wild type controls (Fig. 3C, D), indicating that the oogenesis was largely normal in *taz* mutants.

**Fig. 3.**
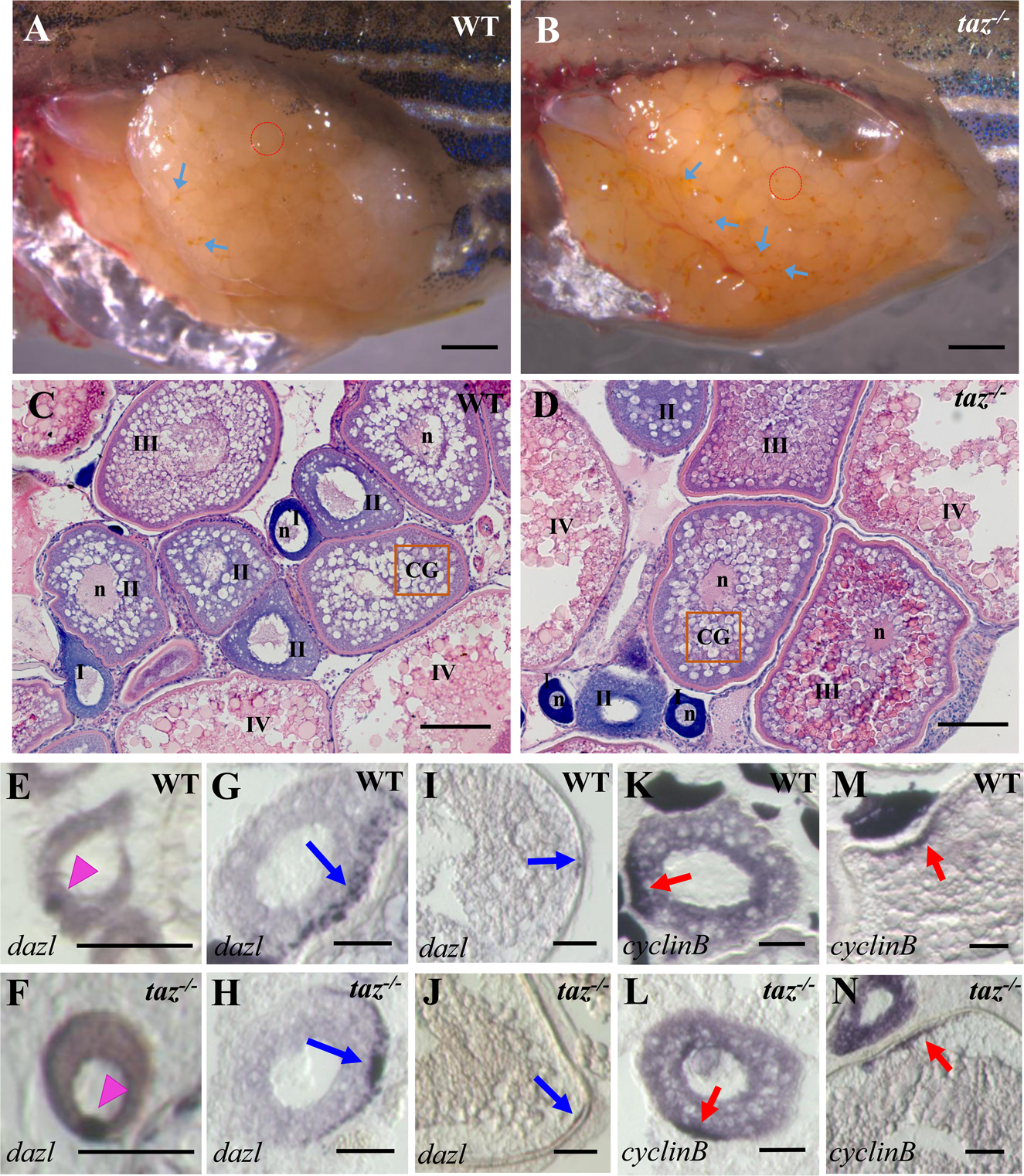
Oogenesis and oocyte polarity is normal in *taz^-/-^* zebrafish females. (**A-B**) Lateral views of adult wild type and *taz^-/-^* ovaries. Oocytes in *taz* mutant ovary (A) are similar to wild type (B) in numbers and size, but we observed more yellow particles (blue arrows) in *taz^-/-^* ovary than in wild type. Red circles outline single oocyte at about IV stage; (**C-D**) Haematoxylin and eosin (HE) stained ovary sections show no apparent morphological difference in oogenesis between *taz^-/-^* (C) and wild type female (D). Oocytes stage I, II, III IV are marked on pictures; n, nucleus; orange box, CG (cortical granules) (**E-N**) *In situ* hybridization to detect oocyte polarity markers. The animal-vegetal polarity is established in *taz^-/-^* oocytes as in wild type oocytes. The Balbiani body, and subsequently, vegetal pole labeled by expression of *dazl* in wild type and *taz^-/-^* primary (E, F), stage II (G, H) and stage III (I, J) oocytes. The animal pole is marked by *cyclinB* transcripts in wild type and *taz^-/-^* oocytes at stage II (K, L) and stage III (M, N). pink arrowhead, Balbiani body; blue arrow, vegetal pole; red arrow, animal pole. Scale bar, 1mm in A-B, 100μm in C-D, and 50μm in E-N.

The establishment of animal-vegetal polarity in oocytes is a key event during oogenesis, and determines the formation of two major embryonic axes, the dorsal-ventral and left-right axis, in vertebrates [29]. Therefore, we examined if the failure of *taz* mutant oocytes to develop was due to defects in animal-vegetal polarity. The Balbiani body (Bb) is the earliest vegetal structure in stage I oocytes, and can be marked by the expression of *dazl* transcripts [30]. In stage I oocytes of *taz* mutants, the Balbiani body had similar animal-vegetal polarity as in wild type oocytes (Fig. 3E, F). Furthermore, we found that *dazl* transcripts were normally located in the vegetal poles in *taz* mutants oocytes through all stages of oogenesis (Fig. 3G-J), and this was confirmed by the distribution of another vegetally expressed transcript, *bruno-like* (*brl*) (Data not shown). The localization of *cyclinB* and *pou2* transcripts, which are animal pole markers, were also similar in *taz* mutant and wild type oocytes (Fig. 3K-N, Data not shown), indicating that the animal polarity was normally established. Taking together, we conclude that *taz* is not necessary to establish animal-vegetal polarity during oogenesis in zebrafish.

### 4. *Taz* is required for micropyle formation in oocytes

Since the oogenesis seemed normal in *taz* mutant ovaries, next we checked if fertilization was normal in mutant eggs. In teleost eggs, the micropyle is a narrow canal for sperm entry through the chorion during fertilization. Unlike a single micropyle on the chorion over the animal pole of wild type fertilized egg, egg water activated egg and stage V oocyte (Fig. 4A-A’, C-C’, E-E’, G-G’), no micropyle was detected in the chorion of *taz* mutant eggs (Fig. 4B, D, F, H). Furthermore, compared with a single obvious cytoplasmic projection from the plasma membrane to the micropyle in wild type chorion (Fig. 4A, E), no protrusion was detected in *taz* mutant eggs (Fig. 4B, F). These observations strongly suggest that *taz* mutant eggs are not fertilized due to the lack of micropyle and sperm likely cannot enter the egg.

**Fig. 4.**
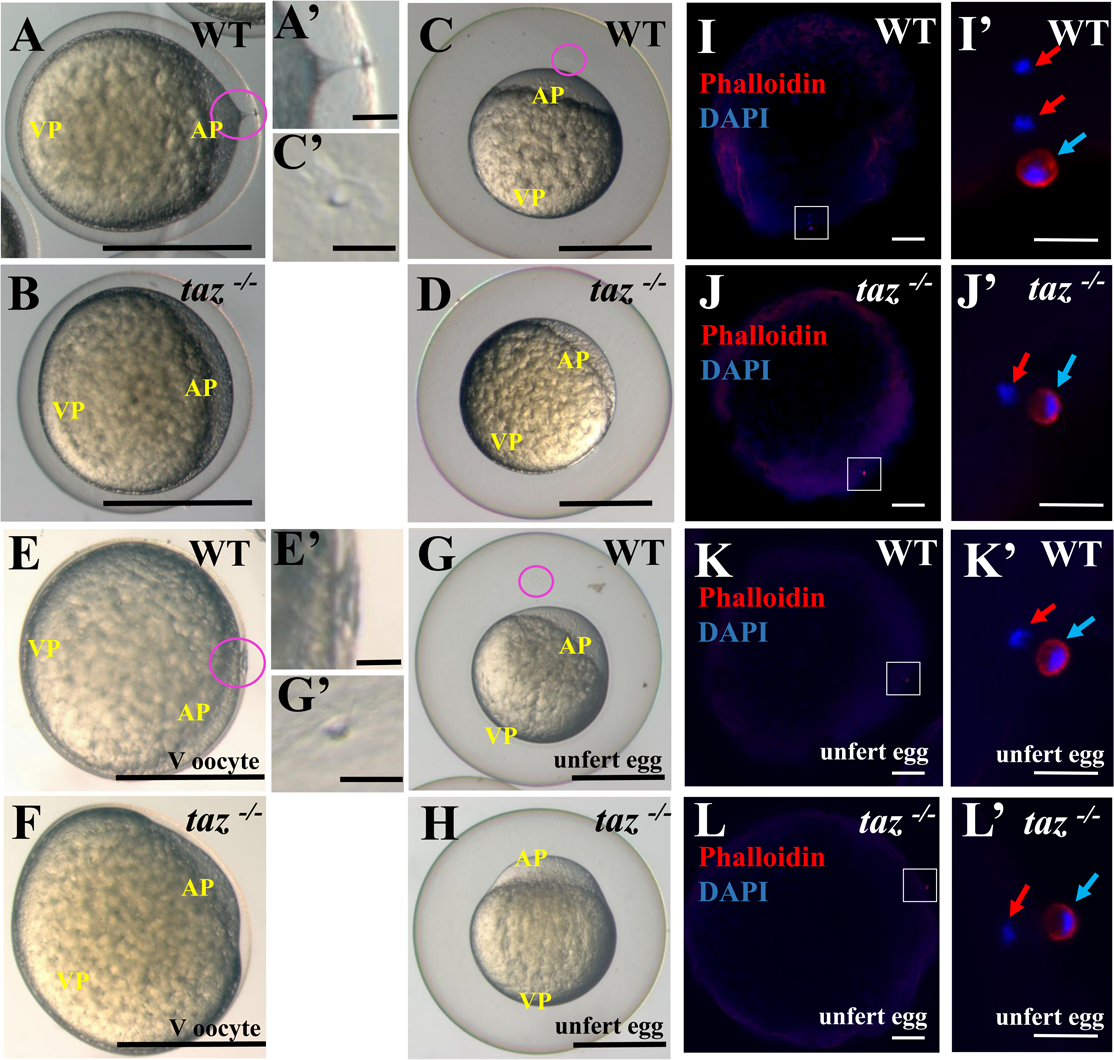
*taz* is essential for formation of the micropyle and fertilization. (**A-H**) A cytoplasmic extension towards the micropyle is observed in wild type eggs activated by fertilization (A, A’), and at 15 minutes post fertilization, a single hole is detected in the chorion above the animal pole of wild type eggs (C, C’). In wild type eggs activated by egg water, a very short extension is seen in stage V wild type oocytes prior to activation by egg water (E, E’) and a hole in the chorion is seen after 15 minutes (G, G’). By contrast, no cytoplasmic projection or micropyle is observed in *taz^-/-^* eggs activated by fertilization (B, D) or egg water (F, H). A’, C’, E’ and G’ are magnified images of the area within the pink circle regions in A, C, E and G. (**I-L’**) Two pronuclei stained by DAPI and one polar body surrounding actin are observed in a fertilized wild type egg at 10 minutes post fertilization (I-I’), whereas one pronucleus and one polar body are detected in an activated *taz^-/-^* egg (J-J’), indicating that *taz^-/-^* egg is not fertilized. One pronucleus and one polar body are observed in unfertilized wild type (K, K’) and *taz^-/-^* (L, L’) eggs. White boxes in E and F denote regions magnified in E’ and F’. red arrow, pronucleus; light blue arrow, polar body; AP, animal pole; VP, vegetal pole. Scale bar, 0.5mm in A-H, 50μm in A’, C’, E’ and G’, 100μm in I-L and 20μm in I’-L’.

Once zebrafish oocytes/eggs are activated (either by egg water or by sperm), the second division of meiosis in oocytes is quickly completed. This is marked by polar body extrusion [31], which is surrounded by Actin. To test if fertilization and meiosis were completed in *taz eggs*, we performed DAPI and Phalloidin staining to detect DNA and the polar body, respectively. When eggs were activated by egg water, both wild type and mutant eggs had one pronucleus, and one Phalloidin marked polar body (Fig. 4K-K’, L-L’), implying that the meiosis could be completed without *taz*. After fertilization with sperm, two pronuclei labelled by DAPI were found in all wild type eggs (Fig. 4I-I’), but only one pronucleus was observed in *taz* mutant eggs (Fig. 4J-J’), indicating that no sperm DNA had entered *taz* mutant eggs.

### 5. Taz is specifically enriched in micropylar and ‘sister’ cells

Oogenesis in *taz* mutant was normal except for the lack of micropyle formation. Besides oocytes, follicle cells are another main group of cells that are essential for oogenesis to progress. In teleost eggs, follicle cells surround oocytes to provide nutrition for their development, and some particular follicle cells specify into unique micropylar cells, which forms one micropyle on each oocyte during stage III oogenesis in zebrafish [32, 33]. To check the status of follicle cells during oogenesis, wild type and *taz* mutant ovaries were sectioned and stained with haematoxylin and eosin (HE). Compared with wild type, in *taz* mutant ovaries, follicle cells around oocytes of all stages had no obvious difference in size, shape and number (Fig. 3C, D). Then we focused on micropylar cells. Remarkably, we found two follicle cells were highly enriched with Taz in early stage III oocytes, while cells around all other stage oocytes examined showed basal levels of Taz expression (Fig. 5A, F). Interestingly, the two follicle cells marked by high Taz expression displayed distinct shapes and spatial positions on the oocyte. One, with an inverted-cone shape, appeared to be inserted into the developing vitelline envelope, and is likely a micropylar cell forming a canal through the envelope (Fig. 5G, I, J). The second cell, which we term as ‘sister’ cell, has a flat shape and tightly covered the base of the cone at early stage III (Fig. 5B, D, E). This cell is not detected from late stage III onwards, and the outcome of this cell is not known. In mutant ovaries the micropylar and ‘sister’ cells were undetectable. The specific enrichment of Taz in the micropylar and ‘sister’ cells implies that Taz might play an indispensable role in the formation of micropyle on zebrafish oocytes.

**Fig. 5.**
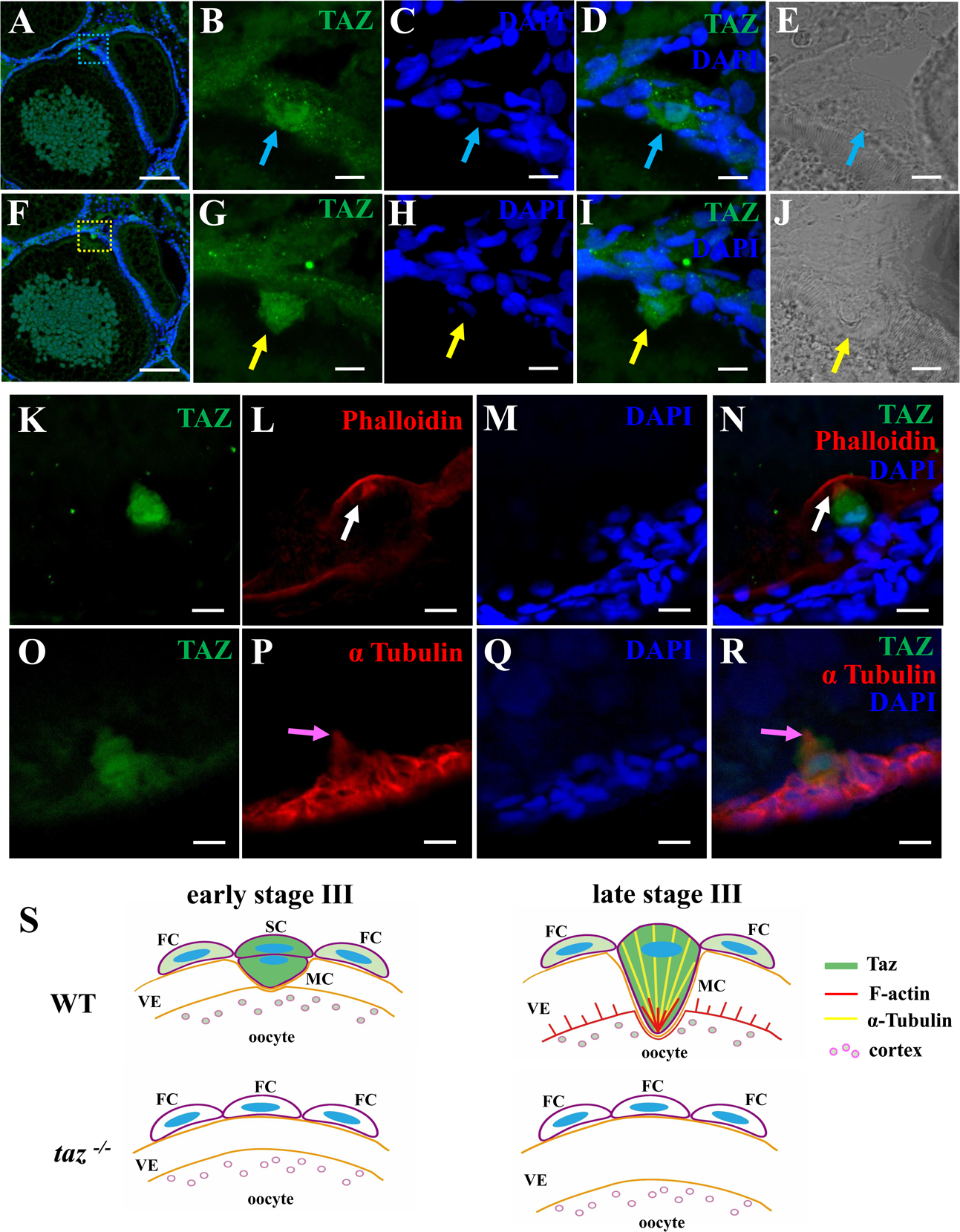
Taz is highly enriched in micropylar cells and co-localizes with the cytoskeleton. (**A-J**) Immunofluorescence of Taz on two consecutive 10μm sections of a wild type ovary. The yellow dotted box (F) shows Taz protein is enriched in a large conical micropylar cell in early stage III oocytes (G), and the blue dotted box (A), shows Taz protein in the sister cell tightly attached to the conical base of micropylar cell (B). An invagination on the vitelline envelope by the protruding micropyle cell is observed (J). B-E are magnified images of the region in the blue dotted box in A, and G-J is the magnified images of region in the yellow dotted box in F. (**K-R**) whole mount immuno-staining of wild type oocyte at late stage III with anti-Taz antibody and cytoskeleton markers. (K-N) F-actin bundles at the leading tip in the cytoplasmic extension of the micropylar cell. (O-R) α-Tubulin is detected in the cytoplasmic extension of micropylar cell. Taz expression overlaps with F-actin and α-tubulin. (**S**) The working model: In early stage III oocytes, two follicle cells (FC), the micropylar and ‘sister’ cells, express Taz. When the conical micropylar cell (MC) depresses the developing vitelline envelope, the ‘sister’ cell (SC) is attached to the base of the micropylar cell. Taz is also expressed in FC and the cortex of oocytes of wild type at a much lower level than in MC and SC. In *taz* mutants, Taz protein is not detected and no MC or SC is formed. At late stage III, the SC disappears, and Taz regulates the cytoskeleton to form a leading tip, composed of F-actin, and Tubulin in the cytoplasm of MC, which might facilitate protrusion into the developing vitelline envelope of oocyte to form a micropyle. This process likely does not happen in *taz* mutants. Samples in A-R were counterstained with DAPI, and oocytes stage I, II, III are marked in A and F. Light blue arrow, the ‘sister’ cell; yellow arrow, micropylar cell; white arrow, the leading tip; pink arrow, cytoplasmic extension. FC, follicle cell; MC, micropylar cell; SC, the ‘sister’ cell; VE, vitelline envelope. Scale bar in A and F, 100μm; in others, 10μm.

### 6. Taz may involve in the formation of the leading tip in the micropylar cell

In many teleosts, formation of the micropyle is thought to require drilling of the vitelline envelope by the micropylar cell, and during this process, the micropylar cell shape undergoes extensive changes [11, 14, 15]. The cytoskeleton might participate in this process. To assess the possible role of Taz in regulating cytoskeletal changes during micropyle formation, we performed co-staining of Taz with Actin and Tubulin in wild type stage III oocytes. We found that Actin filaments constituted an arc leading edge on the vitelline envelope and a leading tip in the micropylar cell (Fig. 5L). Moreover, Taz was found co-localized with actin filament in the leading tip (Fig. 5K, N). Tubulin is also enriched in micropylar cells and overlaps with Taz (Fig. 5O, P, R). These data suggest that Taz may interact with and/or regulate cytoskeletal rearrangements to ensure that the micropylar cell makes a canal through the vitelline envelope to form a functional micropyle.

Taken together, we revealed a unique function of Taz in formation of the micropyle in zebrafish which is summarized in a model (Fig. 5S). In stage III oocytes, Taz, highly expressed in the micropylar cell, interacts with and/or regulates the Actin and Tubulin cytoskeleton, to form protrusions into the vitelline envelope leading to a micropyle. Without Taz, the micropylar cell is not specified, and no micropyle forms in *taz* mutant eggs.

## Discussion

The most interesting finding in this study is that mutations affecting Taz, a key effector of the Hippo signaling pathway lead to loss of a cell required for formation of the sperm entry port on eggs. Our findings identify the first molecular component in the establishment of this unique cell in the zebrafish ovary. Interestingly, Hippo/Taz signaling was found to be deregulated in invasive breast and ovarian cancers [25, 26].

The high expression of Taz in two follicle cells at the animal pole in early stage III oocytes, one of which becomes micropylar cell, is also interesting, and identifies Taz as the first molecular marker for the micropylar cell. The fate of the second cell (‘sister’ cell) and its role in the ovary is currently unclear. It is widely accepted that follicle cells close to animal pole of oocyte contribute towards micropyle formation in many teleosts [12–14]. At this stage, it is hard to distinguish if the high expression of Taz is a cause or consequence of micropylar cell specification from follicle cell. Lineage tracing in wild type and *taz* mutant ovaries, combined with single-cell gene expression profiling can address this question [34–37]. Our work raises interesting questions about how the micropylar cell is specified from a particular follicle cell by Taz expression, and what ensures that there is only one micropylar cell per oocyte.

As an onco-protein, Taz promotes epithelial-mesenchymal transitions (EMT), migration and invasion of human cancer cells, where cell shape changes are prevalent [22]. The micropylar cell changes its shape from a follicular epithelium into a highly polarized cell with a prominent projection, in a process that is overtly similar to EMT in cancer. The micropylar cell bores through the developing vitelline envelope to form a channel, in which the cell shape must change greatly. Both cancer and micropylar cells are dynamic in shape, and therefore, it is reasonable to speculate that Taz works in a similar way in both cells. Our finding that Taz is co-localizes with Actin in the leading tip of the micropylar cell, and also overlaps with cytoplasmic Tubulin, suggests that Taz may interact with and/or regulate the cytoskeleton to drive morphogenesis of the micropylar cell. In support of this possibility, a previous study in the Japanese rice fish *Oryzias latipes*, showed that bundles of microtubules and tonofilaments are formed and elongated in the protruding cytoplasm of the micropylar cell during its penetration of the developing vitelline envelope [15]. Besides the micropylar cell, a ‘sister’ cell at the base of micropylar cells also expresses Taz during mid-oogenesis. However, this cell is not detected later during oogenesis, and its function in micropyle formation and during oogenesis is unknown.

In zebrafish *buc* mutant ovaries, the Balbiani body never forms, leading to an expansion of animal pole-specific gene expression (e.g. *vg1*) and multiple micropyles form in *buc* mutant eggs. Previous studies also found that extra territories of *vg1* transcripts coincide with the locations of ectopic micropylar cells in *buc* mutant oocytes [17]. By contrast, in *taz* mutant oocytes, animal-vegetal polarity is normal and yet, no micropyle forms. Therefore, the polarity of the oocyte alone is insufficient to determine micropyle formation, and additional mechanisms govern micropyle cell fate. Our work identifies a new level of regulation during specification of this cell, and shows that follicle cells at the animal pole induce the formation of the micropyle in a Taz-dependent manner.

## Materials and methods

### Zebrafish strains and embryos collection

Zebrafish (*Danio rerio*) AB^tü^ strain and subsequently generated *taz* mutant lines (*taz^Δ10/Δ10^* and *taz^Δ1Δ1^*) were raised and maintained in the fish facility in accordance with standard procedures [38]. Embryos or oocytes were collected and staged as described [33, 39].

### Genomic DNA extraction

Embryos (or tail fin clips) were lysed in the lysis buffer (10 mM Tris pH 8.2, 50 mM KCl, 0.3% Tween-20, 0.3% Nonidet P40, 0.5 μg/μl Proteinase K (Fermentas)) at 55°C for 14 hours, and then followed by inactivating proteinase K at 94°C for 20 minutes.

### Generation of *taz* mutants by CRISPR/Cas9 system

The target sequence of *taz* gRNA, 5’-GGAGTCTCCCGGGGCT<u>CGG</u>-3’ (PAM site underlined), was located in the exon 1. Zebrafish Cas9 mRNA and the *taz* gRNA was separately synthesized according to the descriptions [40, 41]. After ZCas9 mRNA (300 pg) and *taz* gRNA (50 pg) co-injection into 1-cell stage wild type embryos, and the lysate of about 10 embryos at 24 hpf was used as template for PCR with primers *taz* fw (5’-AGACCTGGACACGGATCTGGA-3’) and *taz* rv (5’-CACTGTATGCACTCCACTAACTGGT-3’). PCR products were sequenced to examine potential indels occurred in *taz* gRNA target region. Embryos co-injected with worked *taz* gRNA and ZCas9 mRNA were raised to adults (F0). F0 fish were screened to identify the founder whose progeny carry the indels in *taz* gene as previously found, and then offspring of identified F0 were raised up. Individual F1 adults were screened again by PCR using genomic DNA from tail fin clips, and indel types in fish were determined by sequencing.

### Genotyping of *taz* mutant

To detect *taz* Δ10 genetype, genotyping primers were designed to amplify specific bands by PCR with a common primer, *taz* fw2 (5’-CGATCGGACGCAGGAGGAACAA-3’), and two individual primers, *taz* wt rv (5’-CGGGTGTGGGAGTGGAGTC-3’) and *taz* Δ10 rv (5’-CGGGTGTGGGAGTGGAGCT-3’). For *taz* Δ1 genetyping, above *taz* fw and *taz* rv primers were utilized to obtain PCR products for sequencing.

### Histology

Wild type and mutant ovaries were dissected out from 8 month old females, and fixed in 4% PFA at 4°C overnight. The fixed tissues were embedded in paraffin and sections were cut at 5-μm thickness using a microtome (Leica). Haematoxylin and eosin staining was performed on them according to the standard protocol.

### *In situ* hybridization

Ovaries were dissected out from adult bodies and fixed in 2% PFA at room temperature for 2 hours. After immersion in 30% sucrose in PBS at 4°C overnight, the ovaries were embedded in O.T.C. compound (Sakura) and frozen in ethanol at −80°C. Frozen tissues were sectioned at 10-μm thickness using a Cryotome (Leica). The serial sections were used for *in situ* hybridization as described previously [42]. The DIG-labeled antisense RNA probes *cyclinB* [32, 43], *pou2* [32, 43], *dazl* [30, 44] and *brl* [45, 46] were applied for marking animal and vegetal poles in oocytes. Whole mount *in situ* hybridization was performed according to method used in a previous report to examine the gene expression pattern of *taz* [47].

### Immunohistochemistry

The frozen sections of ovaries were prepared as above mentioned. The ovary sections and oocytes were performed immunohistochemistry as described previously [42]. Anti-Taz (CST; 1:200) and anti-α-tubulin (Sigma; 1:200) were used as the primary antibodies, and subsequent application of secondary antibodies Alexa Fluor 488 and Alexa Fluor 555 (Life Technology; 1:400) for visualization. 4% BSA in PBS was used for blocking and diluting antibodies. FITC-Phalloidin (Sigma, 1:200) was used to detect F-actin. Before covering with Vectashield (Vector lab), DAPI (Roche) was employed to label the nuclei. Images were acquired on a Zeiss LSM700 confocal microscope.

### Western blot analysis

For preparation of zebrafish protein samples, embryos were homogenized in cold PBS with protease inhibitor (Roche) using syringe (1 ml) and needle (size 23G). The body fragments were collected and heated in whole cell lysis buffer (20 mM NaF, 1 mM DTT, 1 mM EDTA, 0.1 mM Na3VO3, 10 % glycerol, 0.5 % NP40, 280 mM KCl, 20mM Hepes pH7.9) at 100°C for 10 minutes. Then the supernatant of lysate was used for western blot analysis according to the standard protocol [48]. In this study, primary antibodies, anti-Taz (CST, 1:1000) and anti-β tubulin (Thermo, 1:1000) were used, while anti-mouse-IgG-HRP (Thermo, 1:5000) and anti-rabbit-IgG-HRP (Thermo, 1:5000) worked as secondary antibodies.

## Acknowledgment

This work is supported by the National Key Basic Research Program of China 2015CB942800 and 2012CB944502, and The Fundamental Research Funds for the Central Universities XDJK2017A012 and XDJK2014A013. KS was supported by funds from the BBSRC, the Wellcome-Warwick Quantitative Biomedicine Programme, and the Leverhulme Trust.

---

**Fig S1.**
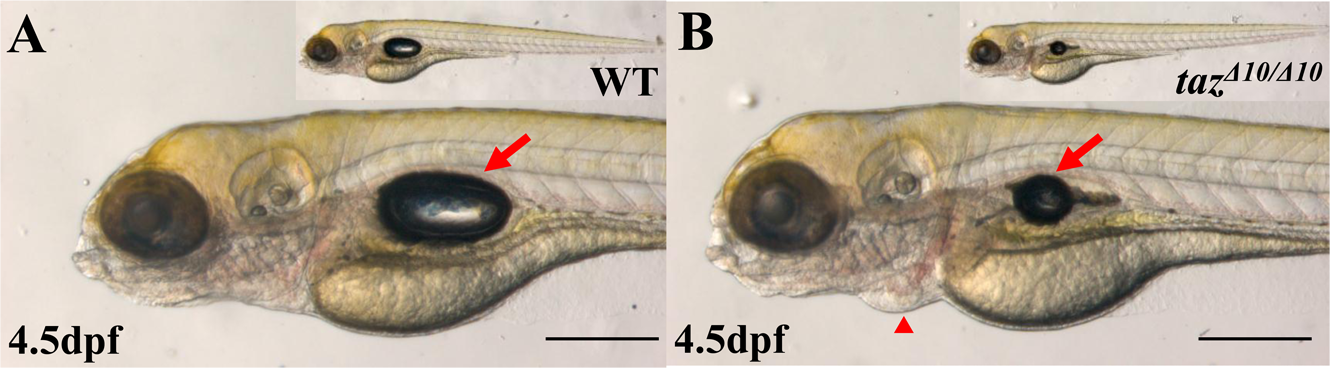
*taz^Δ10/Δ10^* embryos have no lethal defect. (**A, B**) Like wild type (A), *taz^Δ10/Δ10^* embryo is overall normal at 4.5 dpf, except a smaller and inflated swim bladder and mild pericardial edema (B). Red arrow, swim bladder; red arrowhead, pericardium. Scale bar, 500μm.

